# Nonlinear effects of noise on outbreaks of mosquito-borne diseases

**DOI:** 10.1101/2025.08.26.672270

**Authors:** Kyle J.-M. Dahlin, Karin Ebey, John E. Vinson, John M. Drake

## Abstract

Mosquito-borne diseases are a significant and growing public health burden globally. Predictions about the future spread and impact of mosquito-borne disease outbreaks can help inform direct control and prevention measures. However, climate change is expected to increase weather variability, which may shape the future of mosquito-borne disease outbreaks globally. In this study, we sought to determine the effects of demographic and environmental noise (stochasticity) on the duration and size of outbreaks predicted by models of mosquito-borne disease. We developed a demographically and environmentally stochastic Ross-Macdonald model to assess how noise affects the probability of an outbreak, the peak number of cases, and the duration of outbreaks at increasing levels of the basic reproduction number (*R_0_*) and environmental noise strength. Increasing environmental noise lowers the risk of endemic disease from 100% down to almost 0%, but the largest outbreaks occur at intermediate environmental noise levels. In this case, if an outbreak dies out, it ends quickly. With noise present, *R_0_* alone is insufficient to predict definitively whether an outbreak occurs. Surprisingly, our model suggests that increasing environmental noise may reduce the risk of endemic disease and epidemics due to more frequent extreme conditions dramatically affecting mosquito populations.

**Author Summary:** Climate change is expected to cause drastic changes in the spread of mosquito-borne disease outbreaks, both in where they occur and in their size. A key aspect of climate change is an increase in the variability of weather factors, such as rainfall and temperature, factors that also play a crucial role in mosquito survival and reproduction. We created a mathematical model to help us understand how increases in variability might affect mosquito-borne disease outbreaks in the future. Results from our modeling suggest that, depending on current levels, future increases in environmental variability could either increase or decrease the size of future outbreaks. Our work highlights the need to better understand the connections between environmental changes and mosquito biology to inform efforts to forecast and suppress mosquito outbreaks.

## Introduction

Mosquito-borne diseases are a significant and growing public health concern globally, with diseases such as dengue and malaria infecting over half a billion people, causing over half a million deaths annually [1–3]. Vector-borne diseases account for approximately 20% of all newly emerging infectious human diseases [4,5]. The rate of emergence and spread of vector-borne diseases is increasing due to land-use and climate change, meaning that new populations may be exposed to vector-borne diseases with greater frequency [6].

Anthropogenic climate change has already altered and increased the variability in temperature and precipitation and is expected to do so even more in the future [7,8]. Together with these climatic changes, the likelihood of extreme weather events like droughts, heat waves, and extreme rainfall has increased on global and regional scales [9,10]. Climate change and land-use change drive shifts in mosquito species distributions and disease emergence patterns while altering disease transmission dynamics in complicated ways [11]. For example, disease transmission is sensitive to several mosquito and pathogen traits (such as development rate, mortality rate, and biting rate) that are dependent on temperature, often in non-linear ways, leading to disease dynamics that are strongly driven by temperature [12,13]. A major obstacle to anticipating the effects of climate change on disease is poor understanding of the particular form of the non-linear thermal relationship between vector and parasite traits and temperature across species and in different geographic areas [14]. This complexity is evident in empirical studies: a systematic review of the correlation between temperature and dengue transmission found that dengue peaked at intermediate temperatures [15]. In another example, the temporal scale of temperature variation, in this case, daily temperature fluctuations, was found to be a key factor in determining the risk of malaria spread, highlighting the limitations of solely considering average temperature when projecting disease risk [16].

Predicting the spread and impacts of mosquito-borne diseases through mathematical and statistical models can help to inform control and prevention measures. Models have played an instrumental role in improving our understanding of mosquito-borne disease transmission and in determining the optimal targets for control for over a century [17]. Since models are often used to inform public health policies that affect community well-being and mortality rates, modelers strive to make accurate predictions by developing models that can incorporate the biological and social drivers of transmission [18,19].

Environmental variability has long been considered to be an important driver of disease outbreaks and persistence and, more generally, in the stability of ecological networks [20–23]. For example, in some situations where modeling suggests minimal malaria transmission based on mean temperatures, daily temperature variations allow for substantial spread [16]. Demographic stochasticity refers to the inherent randomness in demographic processes like birth, death, and reproduction [24]. Environmental stochasticity refers to unpredictable variation in the environmental factors that regulate demographic processes [24,25]. Past studies have found that ignoring environmental and demographic stochasticity may produce dynamics that do not realistically represent observed patterns in nature, presenting problems for the development of appropriate response strategies [26–28]. The influence that environmental and demographic stochasticity have on the outbreak potential and dynamics of mosquito-borne diseases remains underexplored.

Stochastic models have become more prevalent in public health and ecological research in recent decades as new analytical tools have been developed and improvements to software and hardware have made simulations more practical [29–31]. Like deterministic models, stochastic models can provide predictions of the dynamics of epidemics, including the size and timing of outbreaks [32]. Integrating stochasticity into mathematical models may prove particularly useful for studying the spatial spread of infectious diseases [33]. Stochastic models can provide probabilistic predictions of outbreak occurrence (compared to the binary predictions of deterministic models) as well as the distributions of outbreak sizes and timing by introducing noise that can influence important biological processes for transmission [34,35]. Early warning signals, or statistics derived from noisy epidemic trajectories, can also be used to predict future system states [36–40].

Tools for stochastic modeling of mosquito-borne disease include deterministic models with random parameters [41], stochastic differential equations [27,40], continuous- and discrete-time Markov chain simulation models [42–44], and individual-based models [45–48]. In addition to these mechanistic models, statistical modelers have considered the effects of environmental noise on disease transmission, particularly for vector-borne diseases, using species distribution models [49,50]. Mechanistic models can be used to improve the accuracy of predictions from statistical models by identifying which parameters are most important in determining the system dynamics [51]. While many researchers have considered the effects of stochasticity on mosquito-borne disease outbreaks generally (e.g., its effect on disease emergence), the effect of environmental noise in particular on the characteristics of outbreaks has not been explored.

We modified the Ross-Macdonald mosquito-borne disease model [17,40] to include environmental and demographic noise to explore the dynamics of mosquito-borne diseases in response to stochasticity (Figure 1). By assessing how varying levels of environmental noise affect disease outbreaks, we addressed the following questions: 1) How do demographic and environmental noise affect the probability, intensity, and duration of mosquito-borne disease outbreaks?; and 2) How does the strength of environmental noise affect the probability, intensity, and duration of mosquito-borne disease outbreaks? To explore how the magnitude of environmental noise influences key features of disease dynamics, we focused on three quantities encapsulating the probability and size of outbreaks: the probability of an outbreak of over 100 cases, the peak number of cases, and the duration of the outbreak.

**Figure 1.**
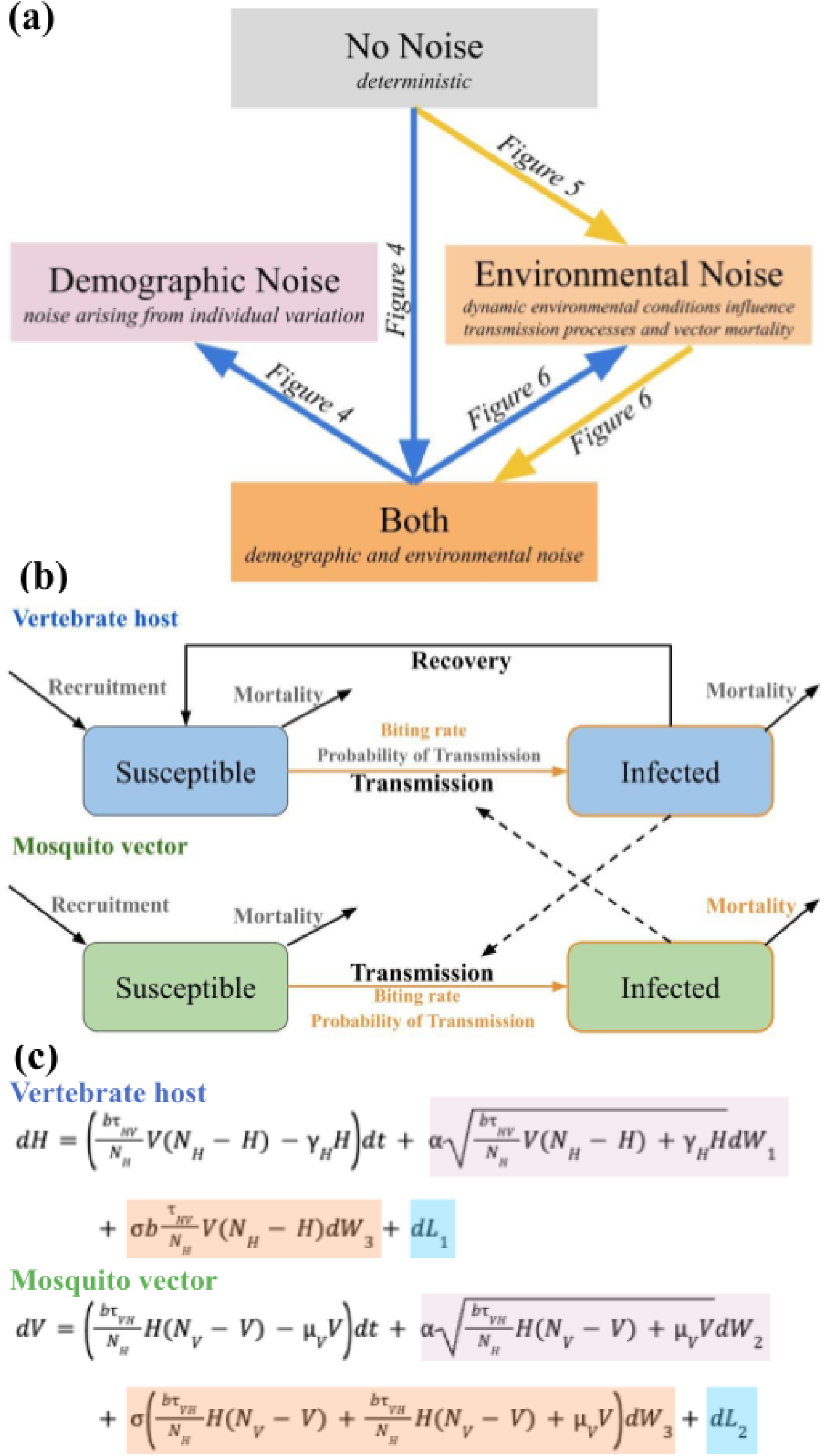
Conceptual diagram. (a) A conceptual diagram of the study highlighting the hierarchical nested layout of the various models. Lines represent the flow of questions explored. (b) Compartmental diagram for the model incorporating demographic and environmental stochasticity. Orange highlights indicate parameters assumed to be impacted by environmental noise. (c) Purple highlights indicate demographic stochasticity, which impacts processes, whereas orange indicates environmental stochasticity, which affects specific model parameters, also colored in orange; 𝜏_𝐻𝑉_is varied so that *R_0_* ranges from 0.75 to 6.5. Total host and vector population sizes are fixed at their base value, while the relative numbers of infected hosts (*H*) and mosquito vectors (*V*) vary over time. The blue highlights represent the reflective processes that keep the model within a biologically meaningful space.

## Results

We begin by describing solutions to the deterministic submodel, then proceed to examine the results of the full model with both types of noise, and finally, to better understand the individual effects of demographic and environmental stochasticity, we consider the demographic noise and environmental noise submodels. We then separately examine the impact of varying the magnitude of environmental noise, 𝜎, on the quantities of interest when demographic stochasticity is included or excluded.

### Deterministic Submodel

Consistent with standard theory, in the deterministic submodel, the probability, intensity and duration of outbreaks depended on *R_0_* in a predictable manner: monotonic increase to an endemic stable state (*R_0_* > 1) or monotonic decrease to pathogen extinction (*R_0_ < 1*) (Figure 2-black lines; Figure 3C, 4C, 5C “Deterministic submodel”). When *R_0_ > 1.01*, an outbreak occurred where at least 1% of the host population was concurrently infected. Intensity increased monotonically with *R_0,_* with approximately 85% of the host population becoming infectious when *R_0_ = 4* (Figure 2; black lines; Figure 4C “Deterministic submodel”).

**Figure 2.**
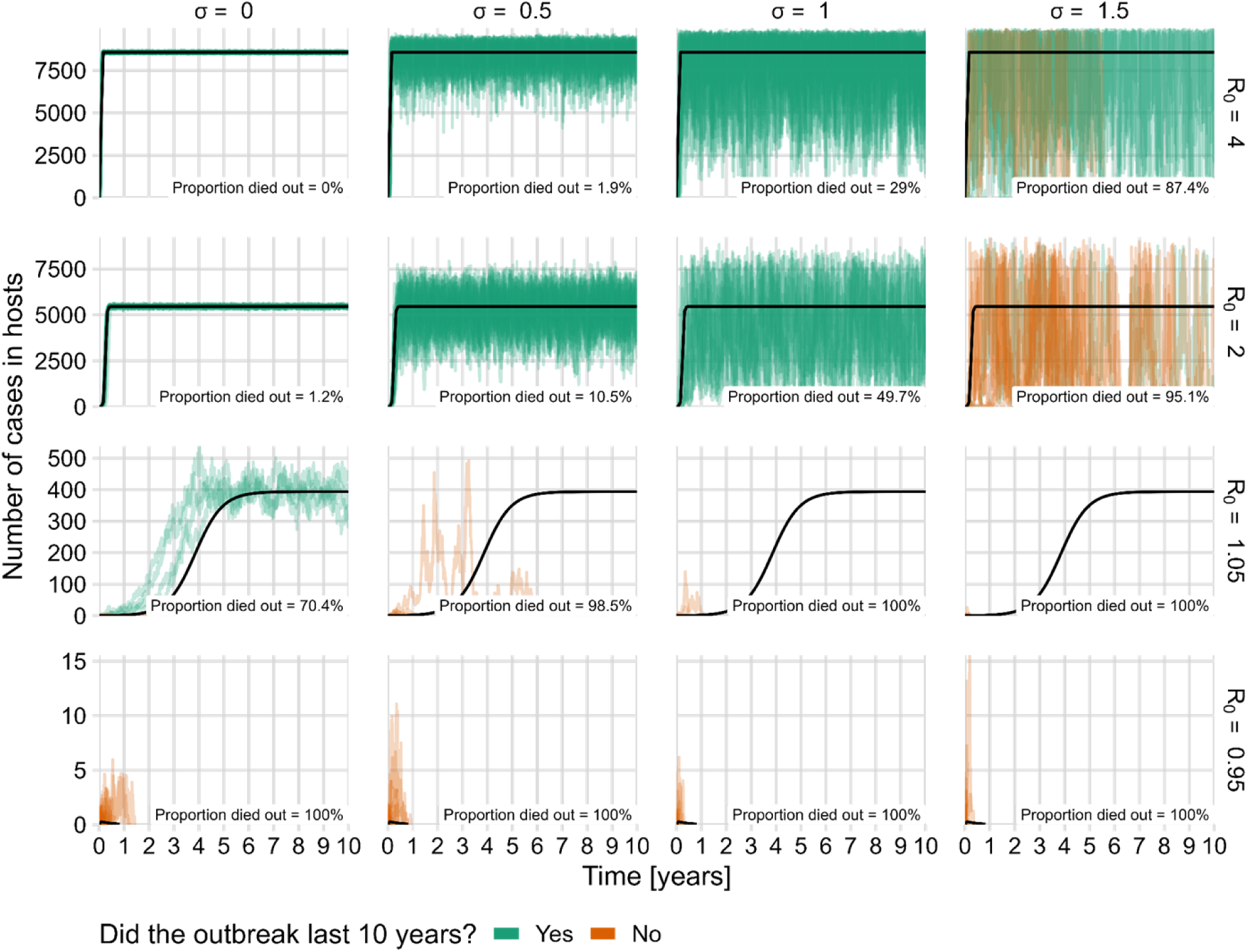
Deterministic model dynamics and 100 sample stochastic trajectories, including both demographic and environmental noise. The number of hosts infected throughout the ten year simulation period is plotted for 16 combinations of four environmental noise levels, 𝜎 (columns) and four *R_0_* values (rows). The black line is the deterministic result for each *R_0_* value. Green lines are stochastic model trajectories where endemic disease was achieved (outbreaks continued for the entire ten year span of simulations) and orange lines are trajectories where the disease died out before the simulations ended. Each panel includes the proportion of simulations in which the disease died out within ten years.

**Figure 3.**
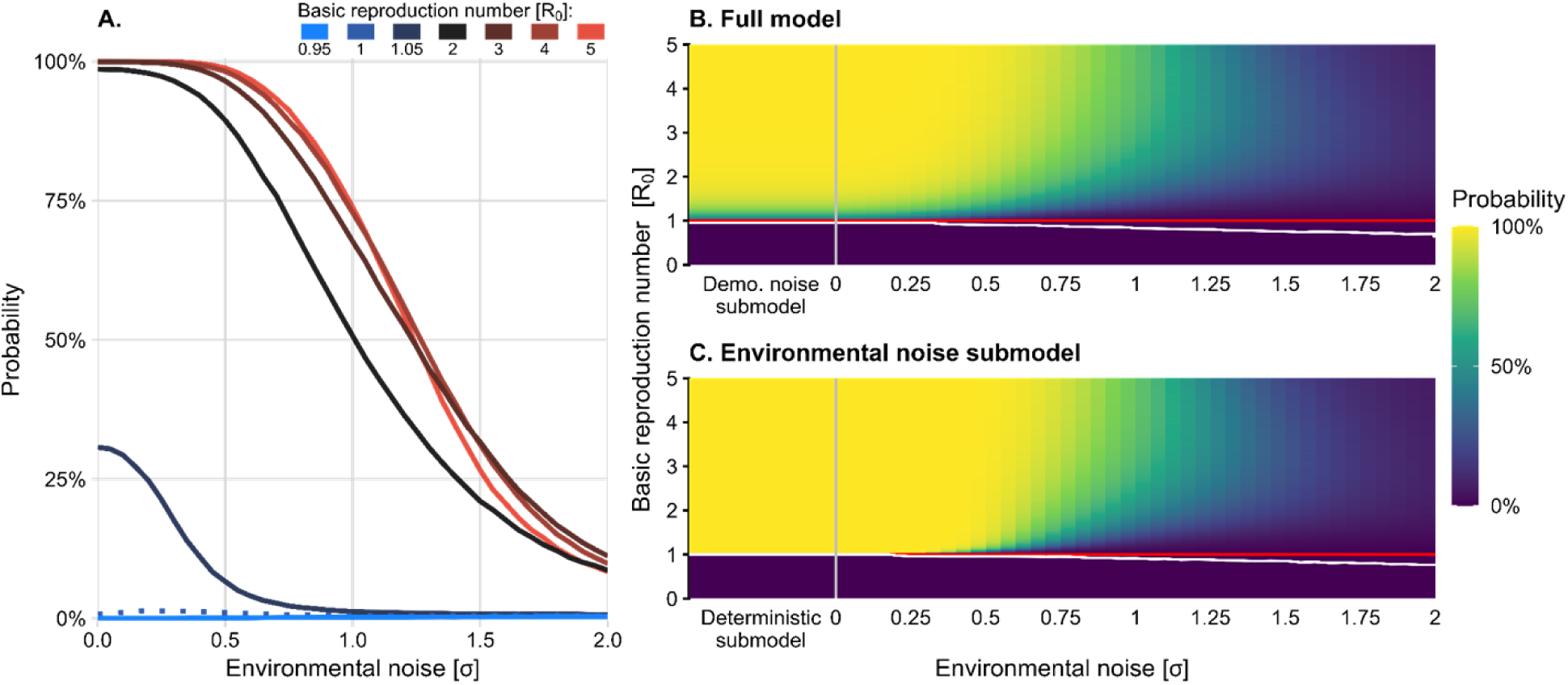
Probability of a large outbreak (mean of the probability of an outbreak exceeding one hundred cases in ten years across 100,000 simulations) (A) The mean of the probability of an outbreak exceeding one hundred cases in ten years across 100,000 simulations as a function of environmental noise strength for seven different *R_0_* values (one 𝑅_0_ < 1 case in blue and the 𝑅_0_ > 1 cases in red shades). (B) The probability of a large outbreak in the full model and (C) in the environmental noise submodel, as both environmental noise strength and the basic reproduction number are varied. The red line indicates the critical threshold of *R_0_=1,* and the white curve indicates the contour above which the probability of an outbreak exceeds 1%. The demographic noise and deterministic submodels are shown to the left of the grey vertical lines in (B) and (C), respectively, where they are equivalent to the cases where 𝜎 equals zero.

**Figure 4.**
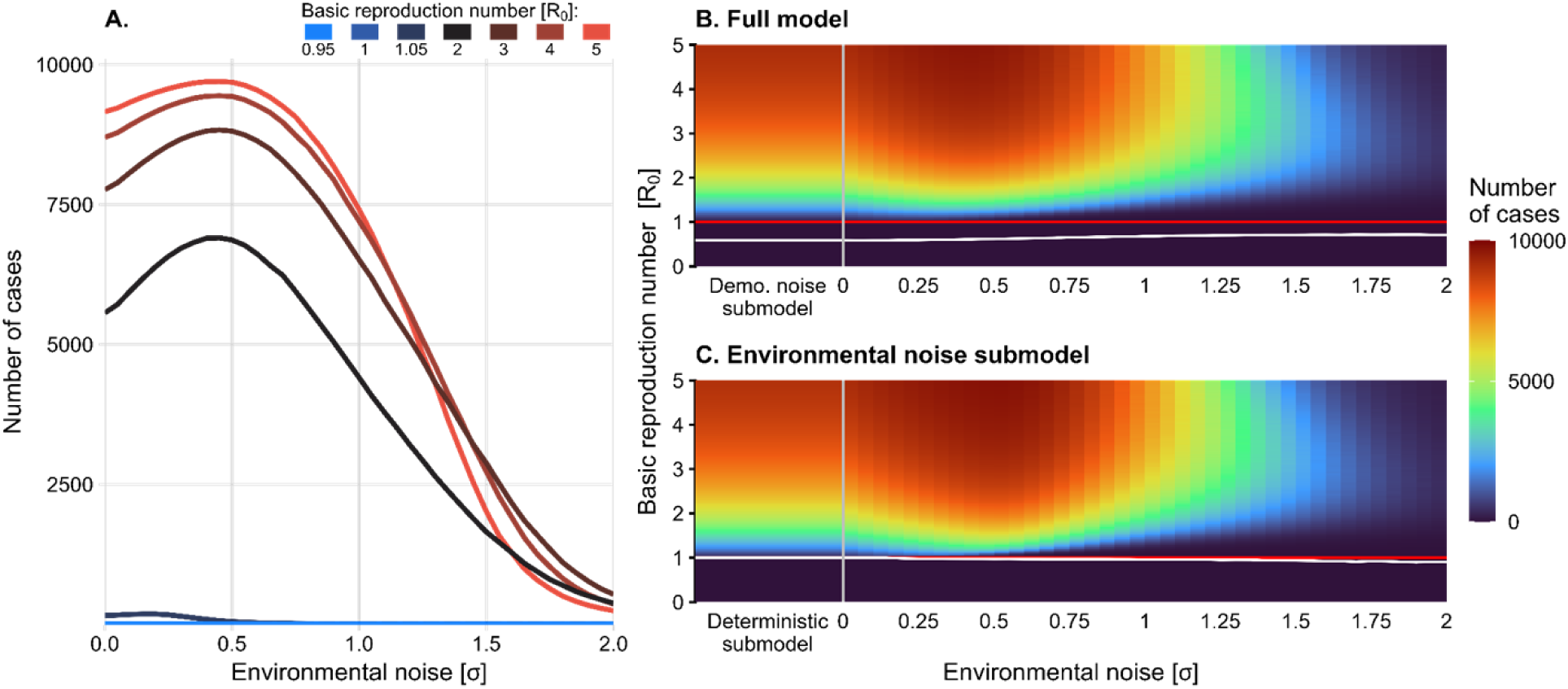
Intensity of outbreaks (mean of the peak number of cases attained in ten years across 100,000 simulations) (A) The mean of the peak number of host cases attained in ten years across 100,000 simulations as a function of environmental noise strength for seven different *R_0_* values (one 𝑅_0_ < 1 case in blue and the 𝑅_0_ > 1 cases in red shades). (B) The intensity of an outbreak in the full model and (C) in the environmental noise submodel, as environmental noise strength and the basic reproduction number, are varied. The red line indicates the critical threshold of *R_0_=1,* and the white curve indicates the contour above which intensity is greater than a single host case. The demographic noise and deterministic submodels are shown to the left of the grey vertical lines in (B) and (C), respectively, where they are equivalent to the cases where 𝜎 equals zero.

### The Addition of Demographic and Environmental Stochasticity

#### Full Model

The addition of demographic and environmental noise led to the possibility of small outbreaks (on the order of 10 cases) lasting up to a year, even when *R_0_* was less than one (Figure 2: bottom row, orange lines; Figure 3A, B). In the deterministic submodel, the probability of a large outbreak is 100% when *R_0_* exceeds 1.01, but demographic and environmental noise reduce this probability, particularly as the strength of environmental noise increases (Figure 3A, B). While the probability of an outbreak often increased with *R_0_* (Figure 3A), the relationship was mediated by 𝜎 (See next section).

Increasing *R_0_* resulted in large oscillations in the number of host cases, typically around the number predicted by the deterministic model (Figure 2). For *R_0_=1.05*, the host cases ranged from close to 0% up to 5% of the population, depending on the value of 𝜎. We observed that at high values of *R_0_*, the entire host population could become infected when 𝜎 was high (Figure 2). Inconsistent with the deterministic model, the average intensity was nonmonotonic with *R_0_*, and peaked when 𝜎 was approximately 0.5 (Figure 4A). Average intensity was non-monotonic in both *R_0_* and 𝜎 (Figure 4B).

For all *R_0_* values, the inclusion of both demographic and environmental noise reduced the average outbreak duration compared to the deterministic submodel (Figure 5A). For low values of 𝜎, higher *R_0_* resulted in longer outbreaks, but when 𝜎 was particularly high, duration dropped precipitously and presented a non-monotonic relationship to *R_0_* (Figure 5B). Threshold-like behavior occurs with duration decreasing from ten years to less than a day as 𝜎 increases from 0.5 to 1.5 for *R_0_* values exceeding one.

**Figure 5.**
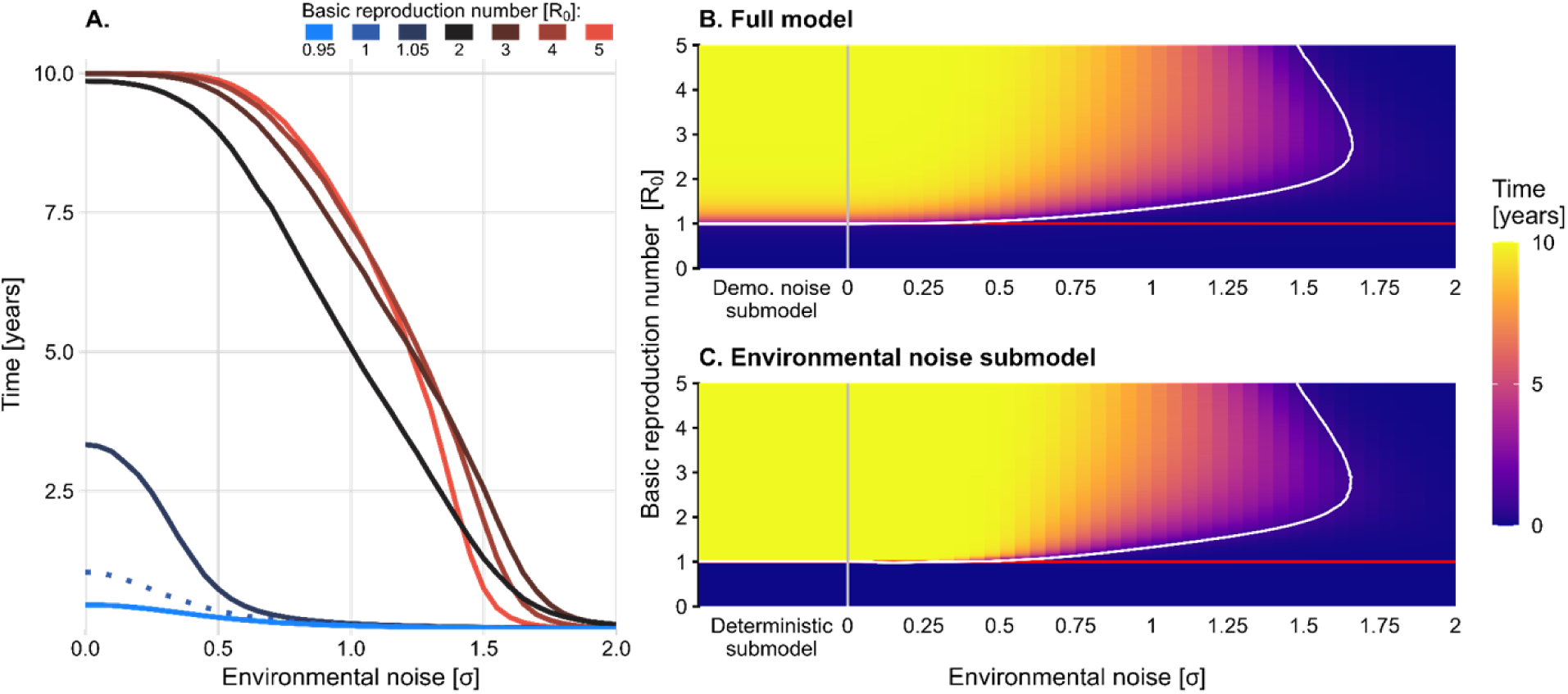
Duration of outbreaks (mean of the final time where there were infections in at least one host and vector across 100,000 simulations) (A) The mean of the final time at which there were infections in at least one host and vector across 100,000 simulations, plotted as a function of environmental noise strength for seven different R_0_ values (one 𝑅_0_ < 1 case in blue and the 𝑅_0_ > 1 cases in red shades). (B) The duration of an outbreak in the full model and (C) in the environmental noise submodel, as environmental noise strength and the basic reproduction number are varied. The red line indicates the critical threshold of R_0_=1, and the white curve indicates the contour above which the duration exceeds one day. The demographic noise (“Demo. noise submodel”) and deterministic submodels are shown to the left of the grey vertical lines in (B) and (C), respectively, where they are equivalent to the cases where 𝜎 equal zero.

#### Demographic noise submodel

For *R_0_* slightly less than one, the inclusion of demographic stochasticity led to more infected hosts compared with the deterministic submodel, but very rarely enough to cause an outbreak of over one hundred cases (Figure 3B, “Demo. noise submodel”). However, when *R_0_* was increased above one, demographic noise reduced the outbreak probability compared to the deterministic submodel (Figure 4B; 6A, “Difference from deterministic submodel”). Outbreak intensity consistently increased with *R_0_*(Figure 4B). Adding demographic stochasticity to the deterministic submodel could cause a slight increase or decrease in outbreak intensity (Figure 6B; “Difference from deterministic submodel”). For all values of *R_0_* > 1, the addition of demographic stochasticity resulted in a shorter outbreak duration compared to the deterministic submodel, particularly for values of *R_0_* slightly greater than one (Figure 6C “Difference from deterministic submodel”). Higher *R_0_* values were associated with longer-lasting outbreaks, on average.

**Figure 6.**
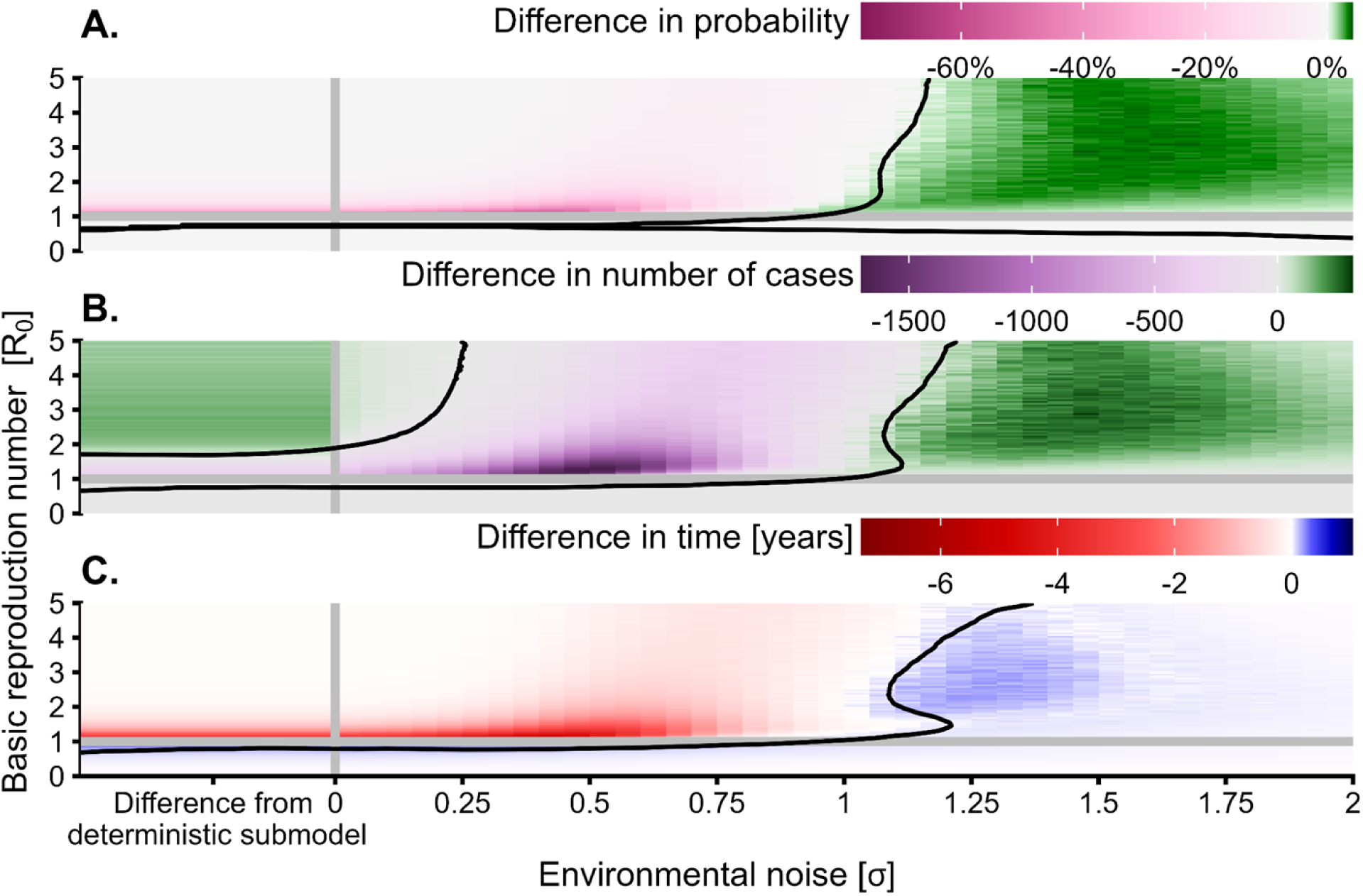
Heatmaps of the difference in probability (A), average intensity (B), and average duration (C) of an outbreak between the full model with both demographic and environmental noise and the environmental noise submodel (i.e., the difference in the values between subplots (B) and (C) in the figures 3, 4, and 5, respectively). The leftmost side of each heatmap indicates the difference between the deterministic submodel and the demographic noise submodel (“Difference from deterministic submodel”). The horizontal grey line indicates the critical threshold of *R_0_=1,* and the black contour lines separate approximate regions of positive and negative differences.

#### Environmental noise submodel

The number of infected hosts in the weakly subcritical environmentally stochastic model (*R_0_* slightly less than one) often exceeded the number in the deterministic submodel, but these outbreaks were rarely large enough to meet our definition (Figure 3C). In general, the probability of an outbreak occurring was higher for larger values of *R_0_*. Environmental noise caused a decrease in the probability of an outbreak for values of *R_0_* greater than one relative to the deterministic submodel, with a substantial increase in this effect at large values of *R_0_*(Figure 6A). In the environmental noise submodel, the average outbreak intensity also had a non-monotonic relationship to *R_0_*, but outbreak intensity increased with *R_0_* at low values of 𝜎 (Figure 4C). The addition of environmental stochasticity to the deterministic submodel resulted in shorter outbreak durations (Figure 5C).

### The Effect of Environmental Noise, 𝜎

#### Environmental noise submodel

The trends observed in the environmental noise submodel, which includes environmental stochasticity but not demographic stochasticity, were similar to the full model. For all values of *R_0_*, increasing environmental noise resulted in lower outbreak probabilities (Figure 3C). When 𝜎 < 1.25, increasing *R_0_*resulted in higher outbreak probabilities, but when 𝜎 > 1.25 outbreak probability was non-monotonic in *R_0_*. Intensity also had a non-monotonic relationship with 𝜎, peaking at intermediate noise levels (Figure 4C). When 𝜎 < 1.25, increasing *R_0_*, on average, resulted in a higher outbreak intensity. But when 𝜎 > 1.25, intensity exhibited a non-monotonic relationship with *R_0_*. Average duration consistently decreased as 𝜎 was increased (Figure 5C). For 𝜎 < 1.25, increasing *R_0_*, on average, resulted in a longer outbreak duration. But when 𝜎 > 1.25, duration again exhibited a non-monotonic trend with respect to *R_0,_* with duration peaking at intermediate values of *R_0_*.

#### Full model

In the full model, outbreak probability decreased with increasing 𝜎 (Figure 3A) when *R_0_*exceeded one. For values of 𝜎 less than approximately 1.25, higher values of *R_0_* led to higher outbreak probabilities. However, for all values of 𝜎 greater than 1.25, as in the environmental noise submodel, there was a non-monotonic relationship between outbreak probability and *R_0_* (Figure 3B). This pattern was especially evident with sharper declines as *R_0_* approaches its maximum value.

The demographic noise submodel exhibited smaller outbreak probabilities than the full model for small values of 𝜎, but slightly greater outbreak probabilities of outbreak for very large values of 𝜎 (Figure 6A). The shift from negative to positive impact of demographic noise on outbreak probability occurs near 𝜎 <u>=</u> <u>1</u> where the overall probability of an outbreak is approximately 50%.

Intensity had a non-monotonic relationship with 𝜎 (Figure 4A). Across all values of *R_0_*, maximum intensity occurred at intermediate values of 𝜎, in the range of 0.4 to 0.6. When 𝜎 less than 1.25, increasing *R_0_* led, on average, to a higher outbreak intensity (Figure 4B). However, when 𝜎 exceeded 1.25, increasing *R_0_* instead caused a decrease in intensity. Comparing the complete model to the environmental noise submodel revealed that the effect of demographic noise on outbreak intensity depended on 𝜎, resulting in a lower outbreak intensity in simulations at low 𝜎 where *R_0_*exceeded one, but slightly increased values when 𝜎 was high (Figure 6B).

Increasing 𝜎 resulted in shorter average outbreak durations (Figure 5A). For 𝜎 < 1.25, increasing *R_0_* resulted in longer average outbreak durations, while at higher values of 𝜎 duration had a non-monotonic relationship to *R_0_* (Figure 5B). The effect of adding demographic noise on outbreak durations was similar to its effect on intensities (Figure 6C).

#### Distributions of intensity and duration across simulations

To better understand how the distributions of intensity and duration vary, we calculated them for select combinations of *R_0_* and 𝜎. At low 𝜎 and *R_0_*, the pathogen immediately went extinct, with no individuals becoming infected (Figure S1, bottom-left plot). However, when the pathogen was able to persist, intensity varied widely. Increasing 𝜎 resulted in more simulations where the pathogen quickly went extinct, but again, outbreaks that persisted had a wide range of intensities (Fig. S1, bottom-right plot). Large values of *R_0_* resulted in a bimodal distribution of intensity where either the pathogen goes extinct quickly with very few individuals infected or persists, and every individual becomes infected (Figure S1, top-left plot). On the other hand, increasing 𝜎 led to an increase in the number of simulations where the pathogen went extinct, but for those short simulations, the intensity was commonly very high (Figure S1, top-right plot).

The reduction in the average outbreak duration with 𝜎 was caused mainly by simulations where the pathogen went extinct nearly instantly (Figure S1). At lower *R_0_*, the pathogen often went extinct immediately, but in some simulations, the duration spanned from zero to ten years. With low 𝜎 and high *R_0_*, the pathogen extinction occurred within days or persisted for ten years. For lower *R_0_* and high 𝜎, the pathogen often became established in the population, but outbreaks occurred for a relatively short duration and none persisted for the entire ten-year period (Figure S1, bottom-right plot). At high *R_0_* and 𝜎, outbreaks could last for brief periods, as the pathogen went extinct immediately. Still, there was potential for the pathogen to persist the entire 10-year period (Figure S1, upper-right plot). In this case, simulations had intense outbreaks (a high number of host infections) but short durations (lasting less than 2.5 years). This highlights that the mean values of stochastic trajectories does not provide a complete picture of the possible dynamics, as some of these trends show bimodality or do not showcase the potentially important “extreme” cases (e.g., short but intense outbreaks).

## Discussion

Our results indicate that the effects of demographic and environmental stochasticity on three key measures of an outbreak (probability, intensity, and duration) are nonlinear and a sound understanding of how stochasticities influence outbreaks is crucial for implementing appropriate responses. The presence of demographic and environmental stochasticity, alone or together, can reduce the probability and duration of outbreaks compared to models with a single source of stochasticity or none, with further decreases in these quantities observed as environmental noise increases. With our stochastic model describing mosquito-borne pathogen transmission, we found that demographic and environmental stochasticity interact to create conditions unfavorable for outbreak persistence, even when *R_0_ > 1*. It is intuitive that demographic stochasticity can reduce outbreak persistence because transmission relies on sufficient interactions between two populations, both of which may be negatively affected by demographic stochasticity. But the mechanism by which environmental stochasticity can drive decreases in outbreak persistence is less clear.

Our finding of the non-monotonic relationship between environmental noise and average outbreak intensity is consistent with observed nonlinear responses between the environment (e.g., temperature) and biological processes for vector-borne parasite transmission (e.g., biting rate) [12]. Daily temperature fluctuations have been shown to control the transmission dynamics of vector-borne parasites, such as the longevity of mosquitoes infected with dengue virus [52,53], viral loads of West Nile virus [54], and several rates associated with malaria transmission [55]. However, others have found that daily temperature variation can have little to no effect on viral load in vectors [56] or infectivity and susceptibility [54]. Our model assumed that daily environmental fluctuations affect the vector biting rate, mortality rate, and the ability of mosquitoes to pass the pathogen on to hosts, all of which may be non-linearly correlated with environmental variables like temperature, rainfall, and humidity [57].

Our results shed light on the potential implications of more extreme environmental variation on outbreak intensity. Outbreaks could become more intense with increased environmental variation, but only if current levels of environmental variation are sufficiently low. Generally, researchers expect that increased environmental fluctuations will lead to more outbreaks [6]. However, counterintuitively, the opposite effect occurs at higher levels of environmental variation, where the environment occasionally acts strongly to increase mosquito mortality or decrease transmission competency, leading to pathogen extinction. It is important to note that in our model, 𝜎 is a simplified composite variable representing the total effect of environmental variation on key mosquito traits; it does not directly represent the nonlinear effects of environmental factors like temperature or rainfall on these traits. However, if increases in environmental variation are expected to lead to more extreme effects on mosquito traits, then 𝜎 is a reasonable proxy.

The observed shift in the relationships between the outbreak measures and *R_0_* when 𝜎 is large might be explained by early crashes in the infected mosquito population when the number of infected hosts is small. Because (𝐻,𝑉) = (0,0) is an absorbing state of the model, if both forms of noise have a strong negative effect at the start of an outbreak, they can quickly push both infected populations to zero.

In models with demographic and environmental noise, outbreak probability, intensity, and duration generally increased with *R_0_*. This is consistent with the deterministic model, where the equilibrium outbreak intensity depends on *R_0_*. However, we found a non-monotonic relationship between *R_0_* and all three disease outcomes, such that the outbreak probability, intensity, and duration are highest at intermediate *R_0_* values. As described previously, at higher values of *R_0_* and sigma, the pathogen may be more susceptible to stochastic extinction. Previous work has shown that for stochastic models, the probability of an outbreak is defined as 1 ― (1/𝑅_0_)^𝑖^, where 𝑖 is the initial number of infectious individuals [58]. This result can be extended to models with multiple infectious compartments [32]. However, while these stochastic models suggest that increasing *R_0_* increases outbreak probabilities, our findings suggest that environmental stochasticity can disrupt this relationship when *R_0_* is large.

Our detailed study of the distributions of outbreak intensity and duration at various *R_0_* and 𝜎 values revealed that changes in their average values were due to bimodality in their distributions. As 𝜎 was increased, more simulations underwent immediate pathogen die out than simulations with no or low environmental noise, leading to decreases in the average intensity and duration values. Thus, when 𝜎 is sufficiently high, there is a substantial divergence in outbreak outcomes where either the pathogen dies out quickly, infecting very few individuals, or the outbreak persists, so that many individuals become infected. Averages are generally utilized for summarizing unimodal distributions as it provides a standardized value for comparisons among groups assumed to be normally distributed. However as we observed here, the distributions of our quantities of interest were bimodal. These results suggest the public health implications of increasing environmental noise are nuanced. Increasing environmental noise may increase the probability of outbreaks dying out quickly. But if outbreaks do become established in a population, they can be just as severe and last as long as outbreaks at low noise levels. Stochastic models of mosquito-borne disease transmission without environmental stochasticity exhibit early warning signals that predict the end of an epidemic [40]. By analyzing the stochastic trajectories preceding the divergence into outbreak persistence or elimination, future work could investigate whether early warning signals for disease elimination exist when environmental stochasticity is included.

Our goal in our study was to isolate the effects of noise on mosquito-borne disease dynamics from other factors that may influence our metrics of interest. To do so, we started with a straightforward yet traditional model of mosquito-borne pathogen transmission [17]. However, this simple model does not include mosquito life stages, including their aquatic stages that are extremely sensitive to environmental conditions such as precipitation and humidity. Extrinsic incubation periods are also sensitive to temperature and are critical indicators of persistence of some pathogens like malaria [12,59]. A more detailed model might be built by including environmental responses to more vector life history traits, better representing the mechanistic relationships between vector life history and variation in abiotic factors like temperature, humidity, and rainfall. Such work could guide future disease mitigation efforts by identifying key mosquito life history traits associated with transmission while also integrating the impacts of environmental and demographic noise on transmission.

Our study represented environmental stochasticity as a phenomenological process that proportionately impacts three mosquito life-history traits. However, the influence of environmental variables on mosquito and pathogen biological processes is probably non-linear and affects each trait differently, so, for example, large environmental changes could have little effect on one trait while strongly affecting another. Realistically, environmental factors cannot be grouped into a single effect, as the interactions between factors strongly affect mosquito life history traits (e.g., humidity most strongly affects mosquito trait performance at higher temperatures) [57]. An analysis of the type presented here would be computationally infeasible for an entirely mechanistic and stochastic model of mosquito-borne disease transmission that incorporates realistic environmental variation. However, through comparison to the results presented here, such a model could be used to better understand how outbreaks are affected by separate environmental factors and their link to mosquito life history traits.

While environmental variation is understood to influence many ecological processes, in particular those involving insects [60–63], it is not commonly incorporated into models of mosquito-borne disease transmission. Our study yielded nuanced and counterintuitive insights about how demographic and environmental stochasticity determine key features of an outbreak: probability, intensity, and duration. In a world experiencing increases not only in average temperatures but also in the amount of variation of such environmental factors, stochastic modeling can be used to predict the distribution of possible futures, improving our understanding of the effectiveness of disease intervention and prevention measures. If the burden of mosquito-borne disease increases and shifts as our planet’s climate changes, it is crucial that we understand the potential non-intuitive and nonlinear effects of a more variable world on the fate of outbreaks.

## Methods

### Model

We studied a system of stochastic differential equations (SDE), a version of the Ross-Macdonald compartmental model of mosquito-borne disease transmission with demographic stochasticity [17]. The model is based on a system of equations previously formulated and studied by [40], but extended to include environmental stochasticity in key mosquito traits and reflection terms to ensure that the state variables remain in the biologically feasible domain. A key assumption is that the total host and vector populations are at their positive equilibria. As a result, the only values changing are the relative proportions of infected hosts, 𝐻, and vectors, 𝑉:

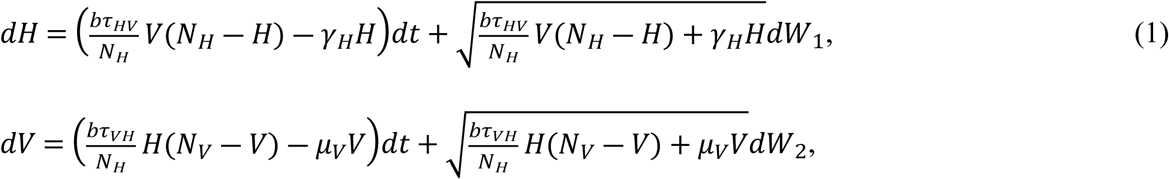

where 𝑊_1_ and 𝑊_2_ are independent Wiener processes. Here, 𝐻 is the number of infectious hosts, 𝑁_𝐻_ is the total number of hosts, 𝑉 is the number of infectious vectors, and 𝑁_𝑉_ is the total number of vectors. The force of infection for hosts is given by 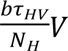 and for vectors by 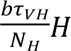. All parameters are defined and described in Table 1.

**Table 1.**
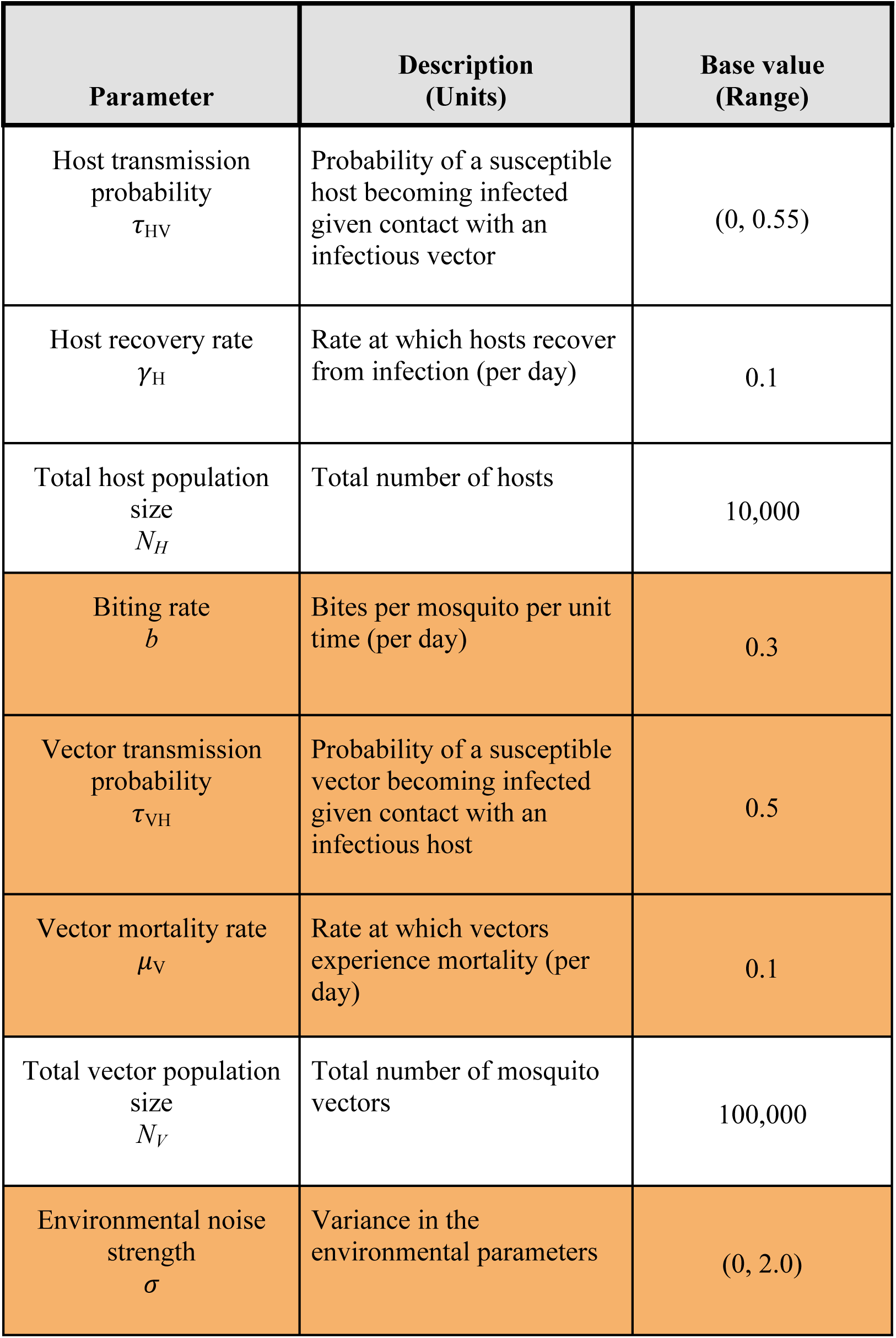
Table of parameters, their definitions, and values. The rows highlighted in orange are parameters that are influenced by environmental noise. The range values for host transmission probability (𝜏_HV_) correspond to *R_0_* values ranging from 0 to 5.

We included environmental noise by perturbing mortality rate 𝜇_𝑉_, biting rate 𝑏, and transmission competency 𝜏_𝑉𝐻_, with an additional Wiener process independent to those for demographic stochasticity. These are the mosquito traits most tightly linked to environmental factors, such as temperature and rainfall [12]. Environmental variation was assumed to proportionally perturb a parameter 𝑔 through the Wiener process described by 𝑔(𝑡)𝛥𝑡 = 𝑔_0_ 𝛥𝑡 + 𝜎𝑔_0_𝜂 𝛥𝑡, where 𝑔_0_ is the value of 𝑔 in the absence of environmental noise, 𝜂 is the standard normal distribution, and 𝜎 denotes the strength of the effect of environmental noise on the parameter. The system of equations in Figure 1b gives the final system of stochastic differential equations. We explored 𝜎 in the range of 0, representing no environmental noise, to a maximum of 2.0, corresponding to the parameters varying up to three times their intrinsic value. The parameter 𝛼 allows us to turn demographic noise on (𝛼 = 1) or off (𝛼 = 0). Environmental stochasticity approximates the effects of random or complex changes in the environment, such as in temperature or rainfall. As an example of how 𝜎 relates to real-world observations, as temperature ranges between 20 and 40°C, the biting rate of a suite of vector-parasite pairs varies between 0.15 and 0.3 [12]. This corresponds to a variation of 33% or 𝜎 = 0.<u>3</u> in our model.

To ensure that solutions of the stochastic model remained in the biologically feasible domain 𝐷 = (0,𝑁_𝐻_ ) × (0,𝑁_𝑉_), which is also the invariant domain of the deterministic model, we modified the equations with reflection terms to arrive at the system of reflected SDEs (2). 𝑑𝐻 =

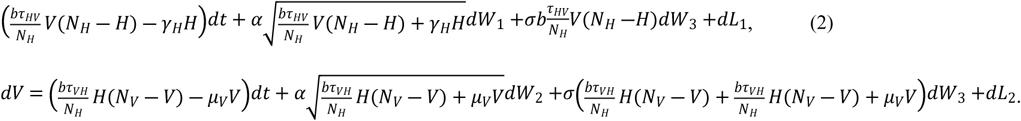

Here 𝑑𝐿_1_ and 𝑑𝐿_2_ are the stochastic processes that reflect the process back into 𝐷. Solutions to system (2) exist and are unique because 𝐷 is bounded [64].

### Parameterization

Without stochasticity, this model admits the basic reproduction number in equation (3) derived using the next-generation method approach [65,66].

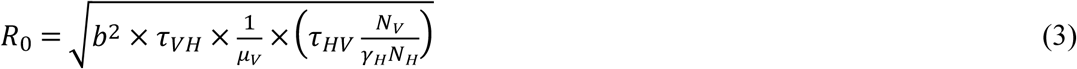

We used the basic reproduction number to determine parameter values such that the deterministic model will lead to persistent transmission (𝑅_0_ > 1) or disease extinction (𝑅_0_ < 1) by fixing all parameters except 𝜏_𝐻𝑉_ then varying 𝜏_𝐻𝑉_ so that 𝑅_0_ varied from 0.05 to 5.0, to include values below and exceeding one, the deterministic threshold for outbreaks.

### Simulations and analyses

#### Quantifying disease dynamics: Quantities of interest

We first conducted an exploratory analysis of model trajectories to better characterize system dynamics over 10 years (Figure 2). Simulations were initialized with ten infectious vectors, and total host and vector populations were set to their maximum values of *N_H_* and *N_V_*, respectively. Solutions to system (2) were approximated using the projection method for the Euler-Maruyama algorithm on the domain 𝐷 with time steps of one-tenth of one day [67]. We observed that even when *R_0_* greatly exceeded one, outbreaks commonly died out at high levels of environmental noise.

Based on these initial observations, we chose to focus our analysis on three quantities: the probability of an outbreak (henceforth, “probability), the peak number of cases (“intensity”), and the duration of the outbreak (“duration”). Understanding these quantities could help decision-makers determine disease risk and prepare for large outbreaks in their communities [68–70]. We defined a realization of the SDE system (2) as an outbreak when the peak number of human cases exceeded 100, that is, when more than 1% of the human population was infectious. Probability was calculated as the proportion of 10,000 realizations that resulted in such an outbreak. Intensity was defined as the maximum number of hosts infected on any one day during the simulation. Duration was defined as the number of days where there was at least one infected host during the simulation or, if the outbreak did not die out, as 10 years, the maximum duration of the simulations. Probability, intensity, and duration were calculated from 10,000 realizations of the SDE system (2).

Because of environmental and demographic stochasticity, each quantity of interest (probability, intensity, or duration ) was a random variable. We calculated each quantity of interest for levels of environmental noise, 𝜎, ranging from 0 to 2.0, and values of *R_0_* from 0.05 to 5, both in increments of 0.025. For each value of 𝜎 and *R_0,_*we obtained a distribution of outcomes. To describe these distributions, we calculated their mean, median, variance, and 25% and 75% quantiles. We primarily report our results using mean values, but this approach assumes a unimodal distribution of our metrics. To investigate whether this assumption held, we explored the qualitative shape of these distributions for selected values of *R_0_* and 𝜎.

#### Hierarchical submodels

We were interested in exploring how demographic and environmental noise can influence the probability, intensity, and duration of outbreaks. System (2) allows for either form of noise to be turned on or off through the parameters 𝛼 and 𝜎. We systematically explored four hierarchical submodels described by system (2) and outlined this process in Figure 1a. The first model included no forms of stochasticity, which we refer to as the “deterministic model” (𝛼 = 0 and 𝜎 = 0). Two models include a single form of stochasticity, the “demographic noise” model (𝛼 = 1, 𝜎 = 0) and the “environmental noise model” (𝛼 = 0, 𝜎 ≠ 0). We also included the complete model that includes “both” forms of noise (𝛼 = 1, 𝜎 ≠ 0). For cases where 𝜎 ≠ 0, we considered values of 𝜎 ranging up to 2.0.

#### Software

While all figures were produced in *R* (version 4.2.3), all simulations were run in the *Julia* language, which can estimate solutions to stochastic differential equations with non-diagonal noise and reflection [71,72]. All code used to produce the results and figures in this manuscript is available in a GitHub repository maintained by the authors and located at the URL https://github.com/DrakeLab/dahlin-noisy-mbds.

## Author Contribution

All authors participated in the design of the study. KD and JV developed preliminary versions of the model. All authors developed the final model, and KD and KE performed the mathematical analysis and computational simulations. All authors drafted the initial manuscript and contributed substantially to revisions. All authors gave final approval for publication and agreed to be held accountable for the content therein.

## Data accessibility

The research uses no data. The R and Julia code used to conduct analyses and generate figures can be accessed at https://github.com/DrakeLab/dahlin-noisy-mbds.

## Acknowledgments

The authors thank the participants of the Population Biology of Infectious Diseases REU at the University of Georgia for valuable feedback.

## Supplementary Material

**Figure S1.**
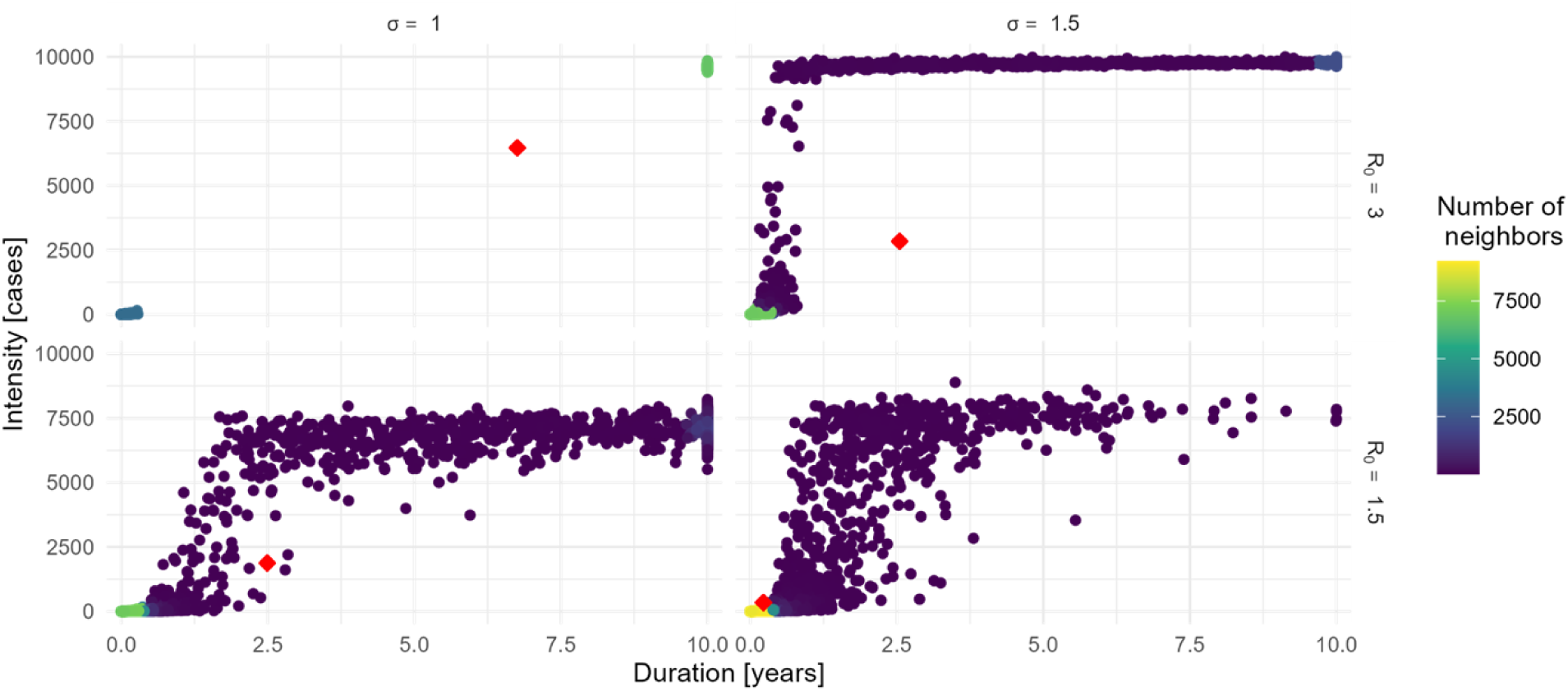
Measurements of the duration and intensity in 10,000 simulation trajectories of the full model. The color of each point indicates the number of its nearest neighbors, signifying the amount of clustering around that point. Each panel is a combination of *R_0_* (rows) and environmental noise strength (𝜎; columns). The red diamond indicates the mean values of duration and intensity.

